# Bovine milk somatic cell transcriptomic response to *Staphylococcus aureus* is dependent on strain genotype

**DOI:** 10.1101/2021.03.04.433893

**Authors:** Dagmara A. Niedziela, Paul Cormican, Gilles Foucras, Finola C. Leonard, Orla M. Keane

## Abstract

**Background:** Mastitis is an economically important disease of dairy cows with Staphylococcus aureus a major cause worldwide. Challenge of Holstein-Friesian cows demonstrated that S. aureus strain MOK124, which belongs to Clonal Complex (CC)151, caused clinical mastitis, while strain MOK023, belonging to CC97, caused mild or subclinical mastitis. The aim of this study was to elucidate the molecular mechanisms of the host immune response utilising a transcriptomic approach. Milk somatic cells were collected from cows infected with either S. aureus MOK023 or MOK124 at 0, 24, 48, 72 and 168 hours post-infection (hpi) and analysed for differentially expressed (DE) genes in response to each strain.

**Results:** In response to MOK023, 1278, 2278, 1986 and 1750 DE genes were found at 24, 48, 72 and 168 hpi, respectively, while 2293, 1979, 1428 and 1544 DE genes were found in response to MOK124 at those time points. Genes involved in milk production (CSN1, CSN10, CSN1S2, CSN2, a-LACTA and PRLR) were downregulated in response to both strains, with a more pronounced decrease in the MOK124 group. Immune response pathways such as NF-κB and TNF signalling were overrepresented in response to both strains at 24 hpi. These immune pathways continued to be overrepresented in the MOK023 group at 48 and 72 hpi, while the Hippo signalling, extracellular matrix interaction (ECM) and tight junction pathways were overrepresented in the MOK124 group between 48 and 168 hpi. Cellular composition analysis demonstrated that a neutrophil response was predominant in response to MOK124, while M1 macrophages were the main milk cell type post-infection in the MOK023 group.

**Conclusions:** A switch from immune response pathways to pathways involved in maintaining the integrity of the epithelial cell layer was observed in the MOK124 group from 48 hpi, which coincided with the occurrence of clinical signs in the infected animals. The higher proportion of M1 macrophages in the MOK023 group and lack of substantial neutrophil recruitment in response to MOK023 may indicate immune evasion by this strain. The results of this study highlight that the somatic cell transcriptomic response to S. aureus is dependent on the genotype of the infecting strain.

## Introduction

Bovine mastitis is a disease of the udder which affects milk yield and quality and causes significant economic losses to the dairy industry (1–3). Mastitis can present as clinical or subclinical. Clinical mastitis presents with overt signs such as clots in milk or swelling of the quarter. Clots in milk are a result of the influx of substantial numbers of immune cells into the udder and are formed by aggregates of leukocytes and blood clotting factors (4). Subclinical mastitis presents with no obvious signs but typically causes a moderate influx of immune cells into the udder, leading to an increase in somatic cell count (SCC) in milk (5). Individual cow SCC of < 100,000 cells/ml is typically considered to be reflective of a healthy mammary gland, while an SCC > 200,000 cells/ml may be indicative of bacterial infection (6).

Staphylococcus aureus is a major cause of bovine mastitis in Ireland (7, 8) and worldwide (9–12) and is also one of the aetiological agents of human mastitis (13, 14). S. aureus is a clonal pathogen which evolves slowly compared to many other bacteria (15). Nevertheless, substantial genetic diversity exists among S. aureus strains, with many lineages identified to date, often adapted to different hosts (16). Four main bovine-adapted lineages have been identified among Irish clinical mastitis isolates: Clonal Complex (CC)71, CC97, ST136 and CC151 (17). CC97 and CC151 are also the most common mastitis-associated S. aureus lineages in Europe (18). CC97 is a predominantly bovine-associated lineage although host jumps into humans with subsequent global spread have been demonstrated (16), while CC151 appears to be a uniquely bovine lineage which has not been found in other ruminants, avian species or in humans (16, 19–21).

During bovine IMI, pathogens enter the mammary gland through the teat canal. The immune response in the mammary gland is presumably initiated by bovine mammary epithelial cells (bMEC) in the alveoli of the udder and by resident macrophages (22). Pathogen-Associated Molecular Patterns such as peptidoglycan and lipoteichoic acid on the surface of Gram-positive bacteria are recognised by Toll-like receptor 2 (TLR2), which stimulates activation of the NF-κB signalling pathway in bMEC as well as macrophages (23, 24). NF-κB and other immune pathways stimulate the production of cytokines and chemokines, which cause inflammation in the udder and stimulate neutrophil recruitment (25). Neutrophils are the first cell type recruited into the udder to engulf the bacteria and attempt to clear the infection (26). Previous studies have reported that S. aureus IMI induces a weak inflammatory response with inhibition of the TLR2 and NF-κB pathways and low levels of pro-inflammatory cytokine production (27–29). However, these studies used a variety of S. aureus strains and the magnitude of the immune response to S. aureus has been reported to be strain-dependent (30). Therefore, the general supposition that S. aureus causes a low immune response in bovine IMI (31) may only be true for specific strains of S. aureus and there is a need to further study the immune response to different strains of S. aureus during IMI.

The transcriptomic response to intra-mammary challenge with S. aureus has been previously studied in bovine gland cistern tissue using microarrays (32, 33) and RNA sequencing (28); however, studies were often short in duration (28, 32) or involved only one time point (33). Characterising the transcriptomic response to S. aureus over several time points would be desirable, since it was previously demonstrated in vitro and in vivo that the response to S. aureus IMI changes over time (30, 34). Furthermore, previous transcriptomic studies utilised only single strains of S. aureus for intra-mammary challenge, and none used a strain belonging to CC151, a globally distributed bovine-adapted lineage. Immune gene expression in bMEC in response to S. aureus differs according to bacterial strain or lineage (25, 34, 35). Intramammary challenge of Holstein-Friesian cows with strains MOK023 (CC97) or MOK124 (CC151) demonstrated that S. aureus strain MOK023 caused mild or subclinical mastitis, while strain MOK124 caused severe clinical mastitis (30). Differences were observed in secretion of cytokines IL-8 and IL-1β into milk between the MOK023 and the MOK124 infected cows, which further suggested that the immune response is strain-specific. Due to the differences observed between the immune response to MOK023 and MOK124 in bMEC in vitro and also in vivo, a further study to elucidate the molecular mechanisms of the host immune response in vivo by utilising a transcriptomic approach was performed. The objective of this study was to identify patterns of gene expression associated with the response to infection with two strains of S. aureus. Expression of cell markers was also examined in order to explore the proportions of immune cells in milk somatic cell samples in response to each strain.

## Results

### Summary of in vivo disease presentation

Milk somatic cells collected during a previously described S. aureus IMI challenge trial were used. The influence of bacterial strain on clinical signs, bacterial load, SCC and the concentration of select cytokines and anti-S. aureus antibodies in milk were previously reported (30). Briefly, cows infected with MOK124 generally developed clinical mastitis, while cows in the MOK023 group developed mild or subclinical mastitis. There was a greater drop in milk yield in the MOK124 group and SCC was significantly higher between 24 and 216 hpi in the MOK124 group compared to the MOK023 group. The MOK124 bacterial load was significantly higher than that of MOK023 at 24 and 31 hpi and subsequently decreased, while MOK023 bacterial load slowly increased and became significantly higher than MOK124 later in the infection (216 to 504 hpi). The levels of interleukin 8 (IL-8) and IL-1β were also significantly higher in milk of cows challenged with MOK124 compared to cows challenged with MOK023 during the first week of infection. Post-infection milk somatic cells were primarily neutrophils, which comprised 80-89 % of somatic cells in the MOK124 group and 41-69 % of somatic cells in the MOK023 group. Infection with MOK124 resulted in a significantly higher proportion of neutrophils in milk than infection with MOK023 (30).

### LukM’

The severity of bovine IMI has been previously associated with secretion of the bi-component leukocidin LukMF’ by S. aureus (36, 37). As the genes encoding this toxin are found in MOK124, but not MOK023, LukM in milk from infected animals was quantified. No LukM’ was found in either MOK023 or MOK124 group milk samples at any time point.

### Sequence and mapping quality

Somatic cells collected pre-infection and at 24, 48, 72 and 168 hpi were subjected to 100 bp stranded paired-end sequencing. On average, 95 % of the bases across all samples had a phred score of ≥ 30 and 98 % had a score of ≥ 20. The mean number of reads per sample was 83.5 M; however, less than 60 M reads were generated from 3 samples from time 0 due to the low number of somatic cells in the samples. The distribution of read counts, phred scores and GC content per sample is available in Supplementary File 1. Quality control, as assessed by FastQC, yielded satisfactory results for all samples. On average, 93 % of reads mapped uniquely to the bovine genome UMD 3.1.93.

### Data exploration

Raw read counts were used for DE gene analysis using DESeq2. After filtering for low counts (sum of reads from all samples < 1) and non-protein coding genes, 17,334 genes were included in further analysis. The expression of the 500 most variable genes was used for principal component analysis (PCA) within the DESeq2 software package. Time 0 (pre-infection) samples clustered separately from infected samples (Fig. 1A). Expression patterns from cows infected with either strain did not separate on the PCA plot at 24 hpi. However, at 48 and 72 hpi, separation between the response to the 2 strains was evident. At 48 hpi MOK124 infected animals’ response had progressed to a state that was maintained until 168 hpi, while the response in MOK023 infected animals progressed more slowly to the same state and samples clustered in distinct groups at 48, 72 and 168 hpi. The difference in progression of gene expression in response to each strain could also be seen in the expression profile of a number of core immune genes across the experiment time course. This was illustrated by the NF-κB subunit 1 (NFKB1) gene, which is a key gene involved in switching on production of immune mediators through the NF-κB signalling pathway. A rapid increase in expression of this gene at 24 hpi was followed by a rapid decrease in MOK124 infected animals. On the other hand, infection with MOK023 caused a more sustained NFKB1 induction (Fig 1B).

**Fig. 1.**
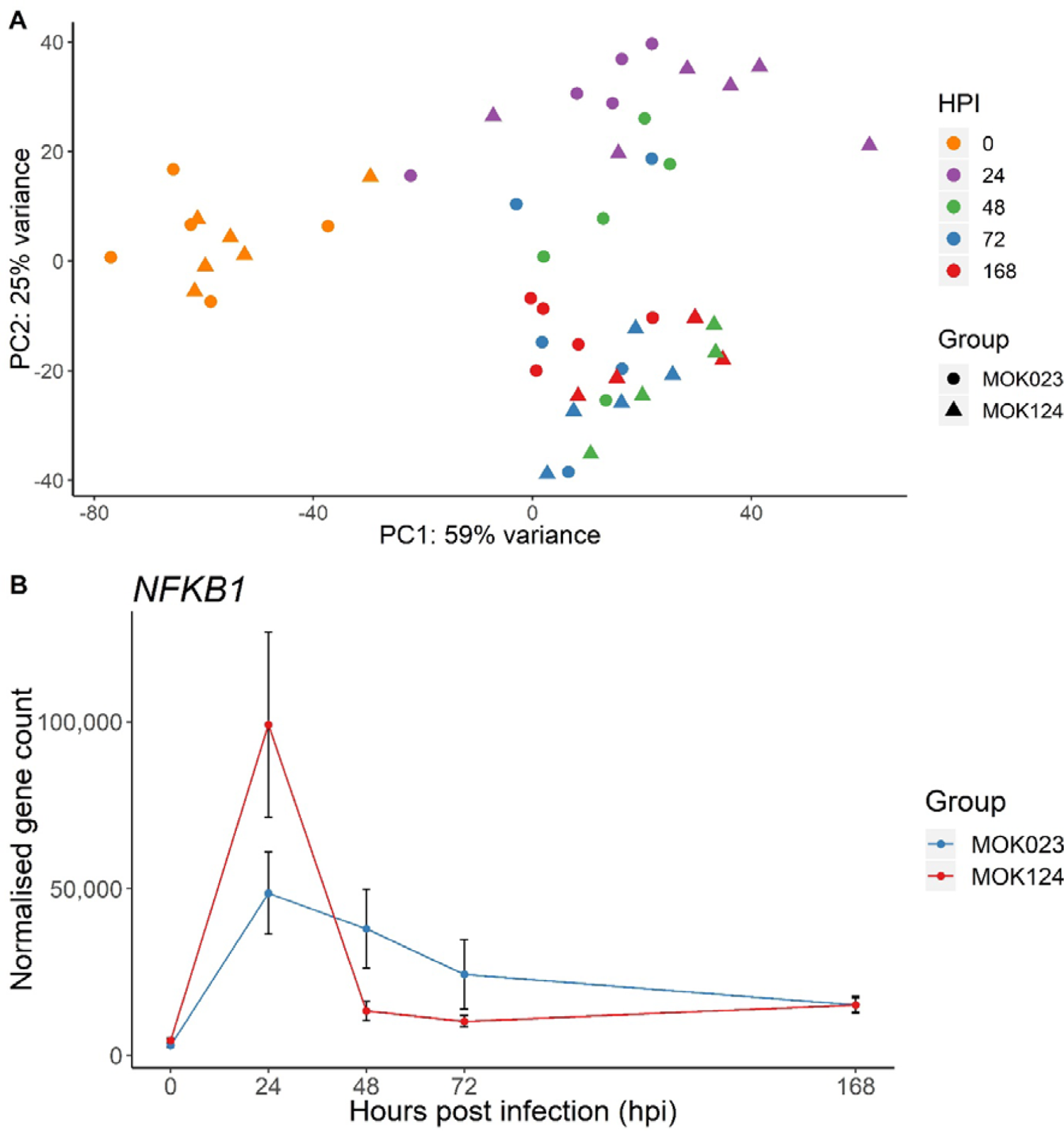
Results of exploration of raw counts data. a PCA plot generated from milk somatic cell gene expression data of the 500 most variable genes following infection with MOK023 (circles) or MOK124 (triangles). Samples from each time point are represented by a different colour: 0 hpi - orange, 24 hpi - purple, 48 hpi - green, 72 hpi - blue, 168 hpi - red. HPI - hours post infection. b Normalised expression (mean ± SEM) of the NF-κB subunit 1 (NFKB1) gene in milk somatic cells in response to infection with MOK023 (blue) or MOK124 (red).

### Differentially expressed genes

To identify highly up and downregulated genes, a log_2_ fold change threshold of 1 (indicating a 2 fold increase or decrease in expression) was applied to the DE gene results in DESeq2 and genes with an adjusted P value < 0.05 were subsequently selected as significant. In response to MOK023, 1278, 2278, 1986 and 1750 significant DE genes were found at 24, 48, 72 and 168 hpi, respectively, while 2293, 1979, 1428 and 1544 significant DE genes were found in response to MOK124 at those time points (Fig. 2A). At 24 hpi the majority of genes with altered expression were downregulated in response to both strains, which could indicate a suppression of normal cell function; while at 72 and 168 hpi, upregulated DE genes predominated in response to both strains. Details of all significant DE genes are available in Supplementary File 2.

**Fig. 2.**
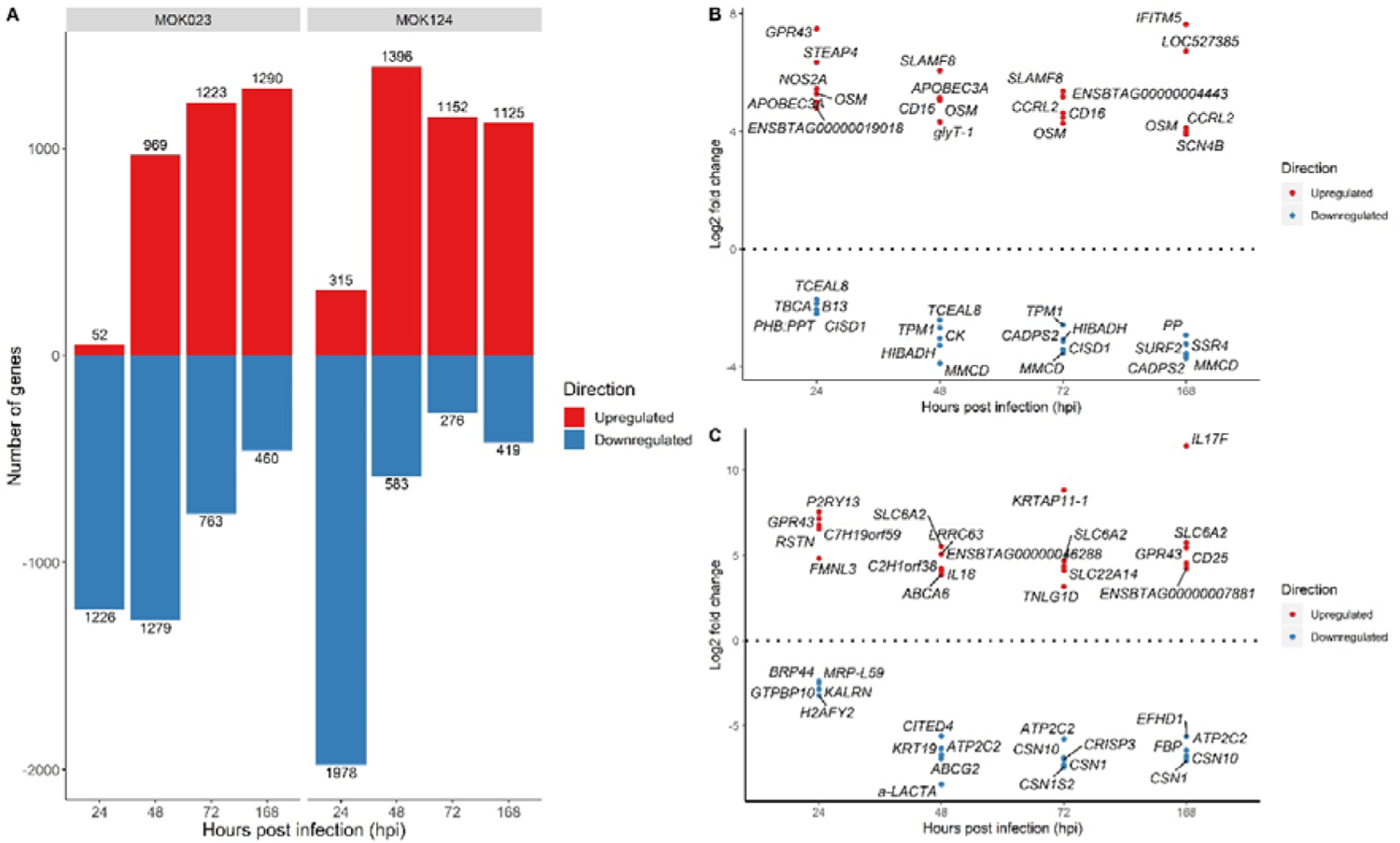
Differentially expressed genes in response to S. aureus MOK023 and MOK124. a The number of significant DE genes (Benjamini-Hochberg adjusted P-value < 0.05, log_2_ fold change threshold > 1) which were up- (red) and downregulated (blue) in milk somatic cells in response to strains MOK023 and MOK124 relative to pre-infection. b Top 5 most significant up- and downregulated DE genes in response to infection with MOK023. c Top 5 most significant up- and downregulated DE genes in response to infection with MOK124. Genes with lowest adjusted P value were selected at each time point. In the case of genes upregulated at 24 hpi in response to infection with MOK023 two genes had the same adjusted P value and so both are included.

Among the most significant upregulated DE genes in response to MOK023, the gene for oncostatin M (OSM) was in the top 5 at all time points post-infection (Fig. 2B). OSM encodes a cytokine involved in inflammation and regulation of production of other cytokines. Oncostatin is also secreted by neutrophils during their rolling in blood vessels in order to enhance neutrophil adhesion to endothelial cells (38). In response to MOK124, the top 5 upregulated genes included those involved in the immune response such as interleukin genes (IL17F, IL18) and interleukin-2 receptor (CD25) gene (Fig. 2C). The top 5 downregulated genes in response to MOK124 included genes encoding milk proteins lactalbumin alpha (a-LACTA), casein alpha subunit 2 (CSN1S2), casein alpha subunit 1 (CSN1) and casein kappa (CSN10).

In order to examine the effect of strain on transcription of major genes involved in milk production, log_2_ fold changes of relevant significantly DE genes were examined (Table 1). The main genes involved in milk production were reviewed by Strucken et al. (2015) (39) and include leptin, lactalbumin, caseins, prolactin and several genes involved in fat transport, energy production and growth. No significant change in expression of the selected milk production genes was observed in MOK023 infected animals at 24 hpi; however, all examined genes with the exception of the gene coding for 1-acylglycerol-3-phosphate O-acyltransferase 6 (AGPAT6) were significantly downregulated at all other time points in this group. In the MOK124 group, downregulation of the expression of milk production genes was already evident by 24 hpi and continued until 168 hpi. Log_2_ fold changes in expression of genes encoding milk-related proteins (a-LACTA, CSN1, CSN10, CSN1S2, CSN2) were 2-3 log lower in the MOK124 group than in the MOK023 group, indicating a more severe reduction of protein synthesis in the first group.

**Table 1.** Expression of genes involved in milk production.

| Gene | MOK023 |  |  |  | MOK124 |  |  |  |
| --- | --- | --- | --- | --- | --- | --- | --- | --- |
|  | 24 hpi | 48 hpi | 72 hpi | 168 hpi | 24 hpi | 48 hpi | 72 hpi | 168 hpi |
| *ABCG2* | N/A | -4.09 | -4.12 | -4.85 | -2.48 | -6.92 | -6.04 | -5.96 |
| *AGPAT6* | N/A | N/A | N/A | N/A | 3.29 | N/A | N/A | N/A |
| α-LACTA | N/A | -4.17 | -4.49 | -5.53 | -2.97 | -8.44 | -7.48 | -7.35 |
| *CSN1* | N/A | -4.09 | -4.29 | -5.27 | -2.80 | -6.72 | -7.31 | -7.04 |
| *CSN10* | N/A | -4.25 | -4.53 | -5.34 | -2.78 | -6.39 | -6.89 | -6.84 |
| *CSN1S2* | N/A | -4.29 | -4.37 | -5.21 | -2.90 | -6.75 | -7.42 | -6.81 |
| *CSN2* | N/A | -4.03 | -4.16 | -5.34 | N/A | -6.71 | -7.22 | -7.13 |
| *GHR* | N/A | -3.18 | -3.47 | -4.16 | N/A | -4.80 | -5.17 | -4.45 |
| *PPARGC1A* | N/A | -4.12 | -3.85 | -4.30 | N/A | -6.93 | -7.06 | -6.93 |
| *PRLR* | N/A | -2.70 | -3.04 | -3.65 | N/A | -4.35 | -3.19 | N/A |

Log_2_ fold changes of significant DE genes only are included (adjusted P-value < 0.05, log_2_ fold change threshold > 1). N/A – gene not significantly DE. Gene names: ABCG2 – ATP binding cassette sub-family G member 2, AGPAT6 - 1-acylglycerol-3-phosphate O-acyltransferase 6, a-LACTA – alpha lactalbumin, CSN1 - casein alpha subunit 1, CSN10 – casein kappa, CSN1S2 - casein alpha subunit 2, CSN2 - casein beta, GHR - growth hormone receptor, PPARGC1A - peroxisome proliferator activated receptor gamma coactivator 1 alpha, PRLR - prolactin receptor. hpi - hours post-infection.

A total of 402 genes in the MOK023 group were DE at all time points post-infection, while 362 DE genes were common in the response to MOK124 at all time points (Fig. 3 A-B). Of these, 130 were shared between MOK023 and MOK124 at all time points and are proposed as strain-independent biomarkers of S. aureus IMI during the first week of infection (Fig. 3C). Among the common genes, 42 were involved in immune response processes.

**Fig. 3.**
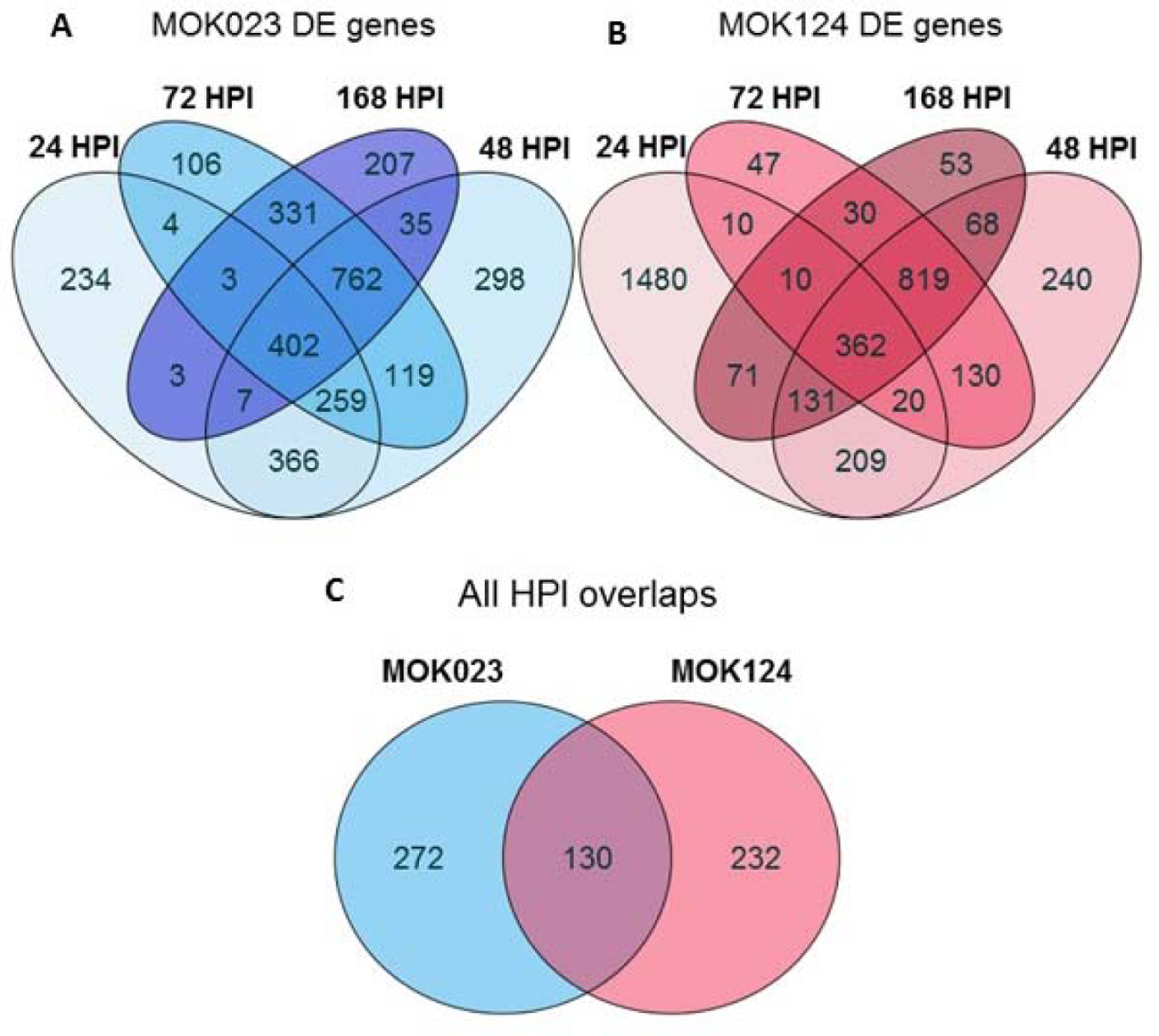
Common DE genes in milk somatic cells across time points. a Common DE genes in response to MOK023. b Common DE genes in response to MOK124. c Overlap of the DE genes common at all time points between the two strains. HPI - hours post infection.

### Pathway analysis KEGG pathways

Pathway analysis of KEGG metabolic processes revealed 67 unique significantly overrepresented pathways in response to MOK023 and 74 significantly overrepresented pathways in response to MOK124. The pathways were enriched at one or more time points. Overall, 84 unique pathways were enriched in response to either strain. The top 10 most significant KEGG pathways at each time point are presented in Fig. 4, while a complete list of pathways at all time points is in Supplementary File 3.

**Fig. 4.**
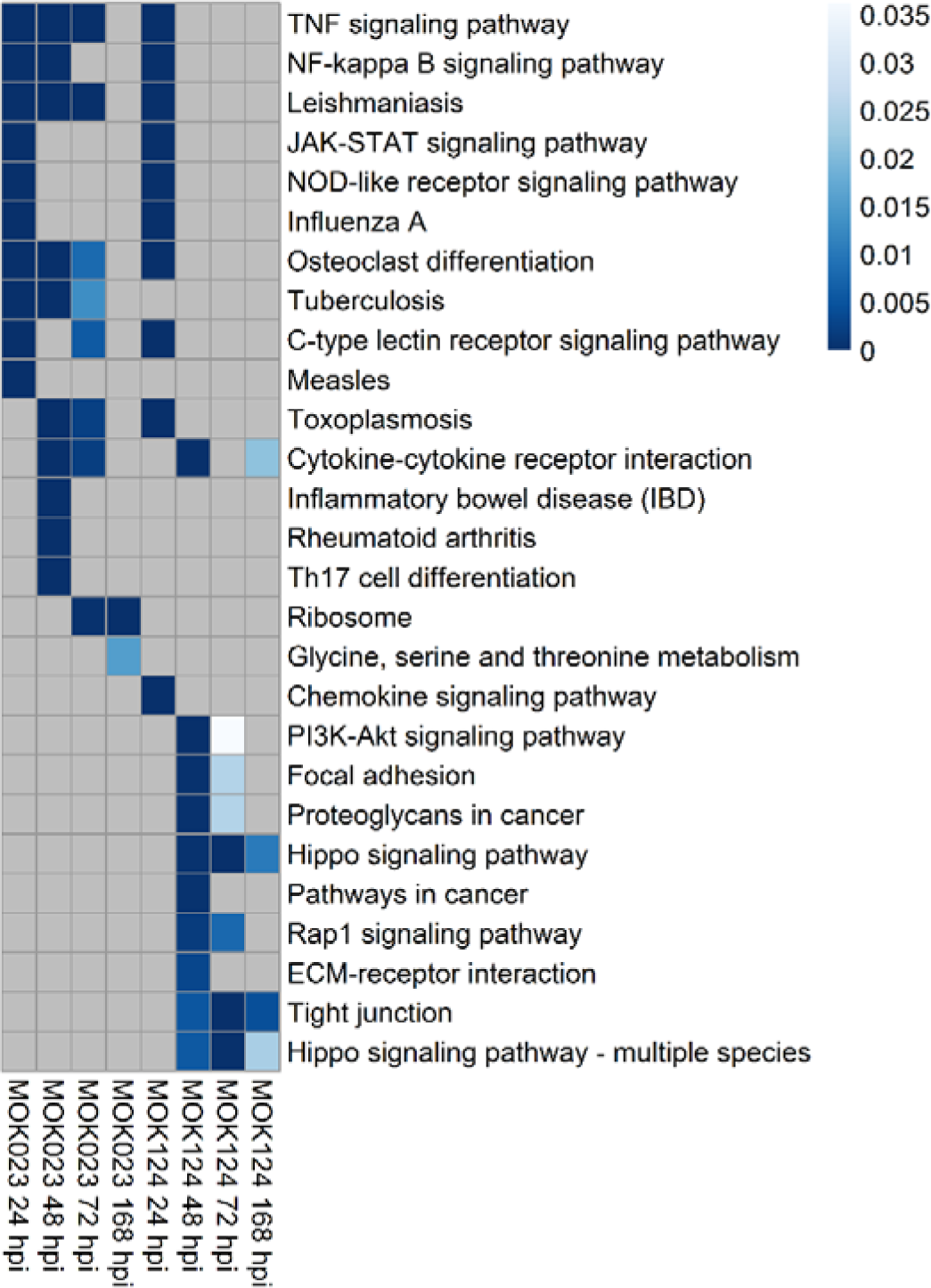
KEGG pathways in response to infection with S. aureus MOK023 or MOK124 over time. Significant (P < 0.05) and unique (filtered for shared genes) are shown. Colour scale represents P value, grey squares represent non-significance. Up to 10 of the most significant pathways at each time point are presented. hpi - hours post infection.

KEGG pathway analysis illustrated that there was a sustained somatic cell immune response to MOK023. At 24 hpi, the top 2 most significant pathways in response to both MOK023 and MOK124 were NF-κB and TNF signalling pathways, both of which are involved in the immune response (Fig. 4). Genes involved in these two pathways displayed similar patterns of expression change in response to both strains at 24 hpi (Fig. 5). Immune response pathways continued to be significantly upregulated in the MOK023 group at 48 and 72 hpi, but not in the MOK124 group, even though mastitis was more severe in the MOK124 group (Fig. 5). Instead, a specific response to MOK124 was evident at later time points, characterised by overrepresentation of the Hippo signalling, extracellular matrix receptor interaction (ECM) and tight junction pathways. The Hippo signalling pathway is involved in regulating cell apoptosis or proliferation (40), while ECM receptor and tight junction pathways both represent groups of genes that influence communication between epithelial cells and maintaining the integrity of the epithelial cell layer (41, 42). When examining genes involved in the Hippo signalling pathway, the majority of genes were downregulated relative to time 0 in response to MOK124 (Fig. 6A).

**Fig. 5.**
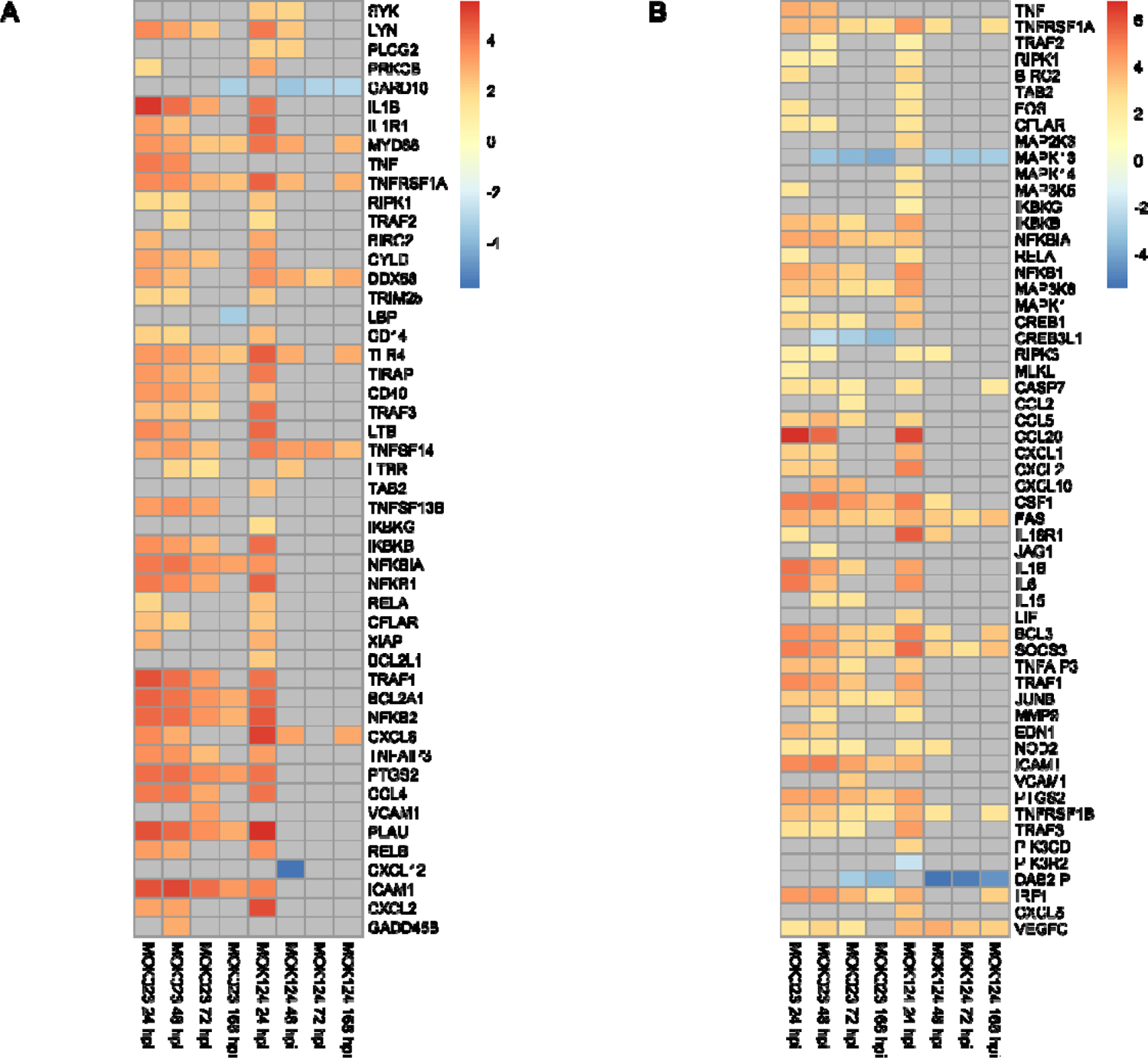
Log_2_fold change of genes involved in selected KEGG pathways in response to MOK023 and MOK124. a NF-κB signalling pathway. b the TNF signalling pathway. Grey tiles show genes that were not significantly DE.

**Fig. 6.**
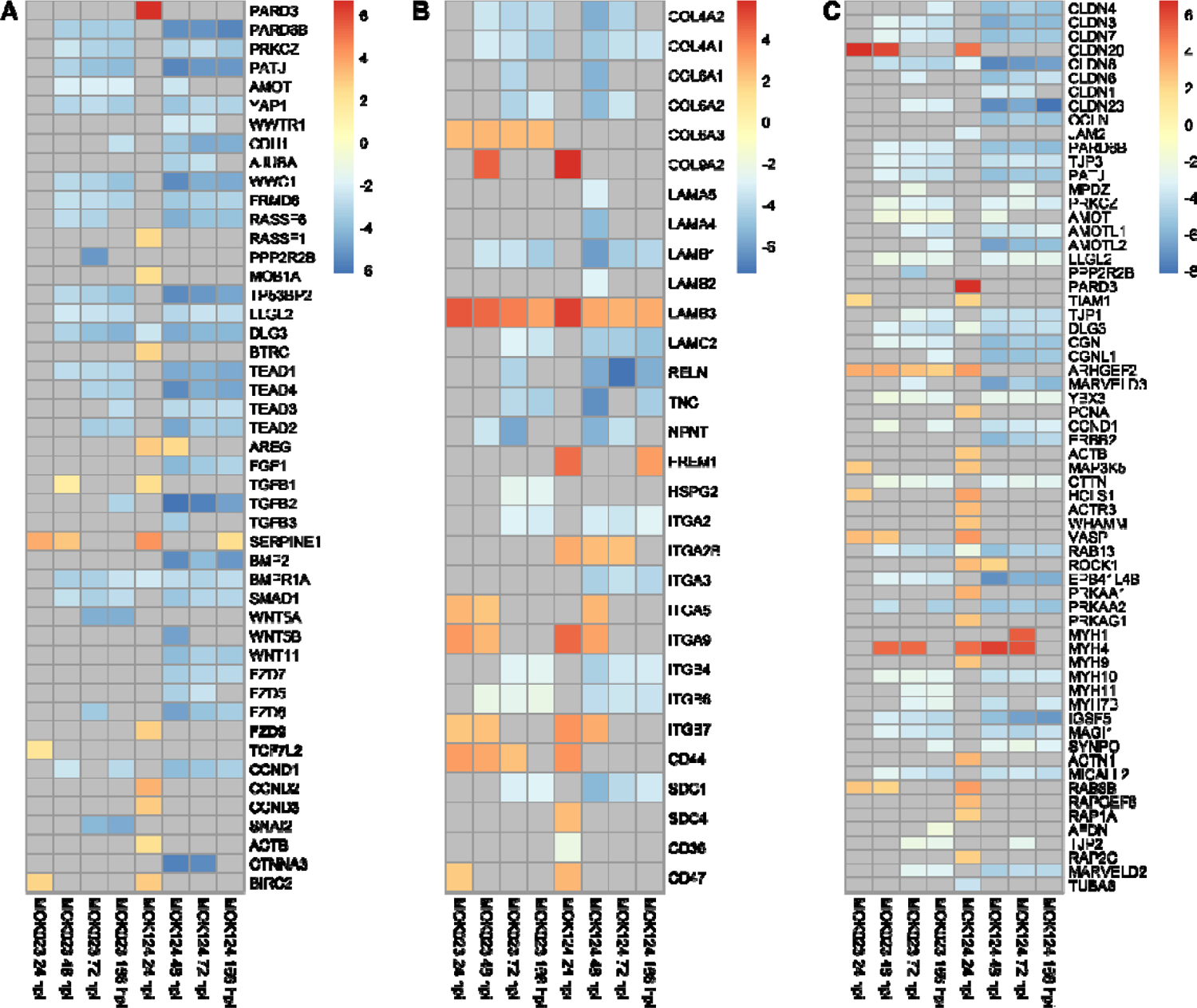
Log_2_fold change of genes involved in selected KEGG pathways in response to MOK023 and MOK124. a Hippo signalling pathway. b ECM receptor interaction pathway. c tight junction pathway. Grey tiles show genes that were not significantly DE.

The ECM receptor interaction pathway was enriched in the MOK124 group at 48 hpi. Among ECM receptor interaction genes, collagen genes COL4A1, COL4A2, COL6A1, COL6A2, COL8A2, COL12A1, COL14A1 and COL18A1 were downregulated in response to MOK124 between 48 and 168 hpi (Fig. 6B). The collagen genes encode proteins that are components of basement membranes. Several integrin genes were upregulated in response to MOK124 at 48 hpi (ITGA5, ITGA9, ITGA2B, ITGB7), while some were also downregulated between 48 and 168 hpi (ITGA2, ITGA3, ITGB4, ITGB6).

For the tight junction pathway, in response to MOK124, most paracellular space genes were downregulated, including genes encoding claudin (CCLN), occludin (OCLN), Marvel domain-containing protein 3 (MARVELD3) and immunoglobulin superfamily member (IGSF5) (Fig. 6C). Several genes from this pathway were upregulated at 24 hpi; however, the pathway was not significantly enriched in the MOK124 group at that time point. Only two genes were upregulated at later time points: Rho associated coiled-coil containing protein kinase 1 gene ROCK1 was upregulated at 48 hpi and myosin heavy chain 1 gene MYH1 was upregulated at 72 hpi, both of which are involved in regulation of actin cytoskeleton. All other genes, involved in cell polarity (PARD6B, TJP3, PATJ, AMOT, PRKCZ, LLG2, TJP1, DLG3), decreased paracellular permeability (TJP1, CGN, CGNL1, CLDN3), reduced cell proliferation (YBX3, CCND1), cell differentiation (YBX3, ERBB2) and tight junction assembly (RAB13, TJP1, CGN) were downregulated between 48 and 168 hpi.

While pathways specific to MOK124 emerged from 48 hpi, the tuberculosis pathway was solely overrepresented in response to MOK023, at 24, 48 and 72 hpi. Notably, among the upregulated genes in this pathway were CTSD, ATP6A1 and LAMP1, which encode cathepsin D, ATPase H+ transporting accessory protein 1 and lysosomal associated membrane protein 1, respectively. All of these proteins are involved in internalisation into macrophages and arrest of phagosome maturation. Other pathways specific to MOK023 were inflammation related pathways, such as inflammatory bowel disease (IBD) pathway and rheumatoid arthritis pathway.

### Gene ontology

Enrichment analysis of GO biological processes revealed 36 different significantly overrepresented terms in response to MOK023 and 36 significantly overrepresented terms in response to MOK124. Overall, 66 different GO terms were enriched in response to either strain. The top 10 enriched terms in response to each strain at each time point are presented in Fig. 7, while a complete list at each time point is attached in Supplementary File 3.

**Fig. 7.**
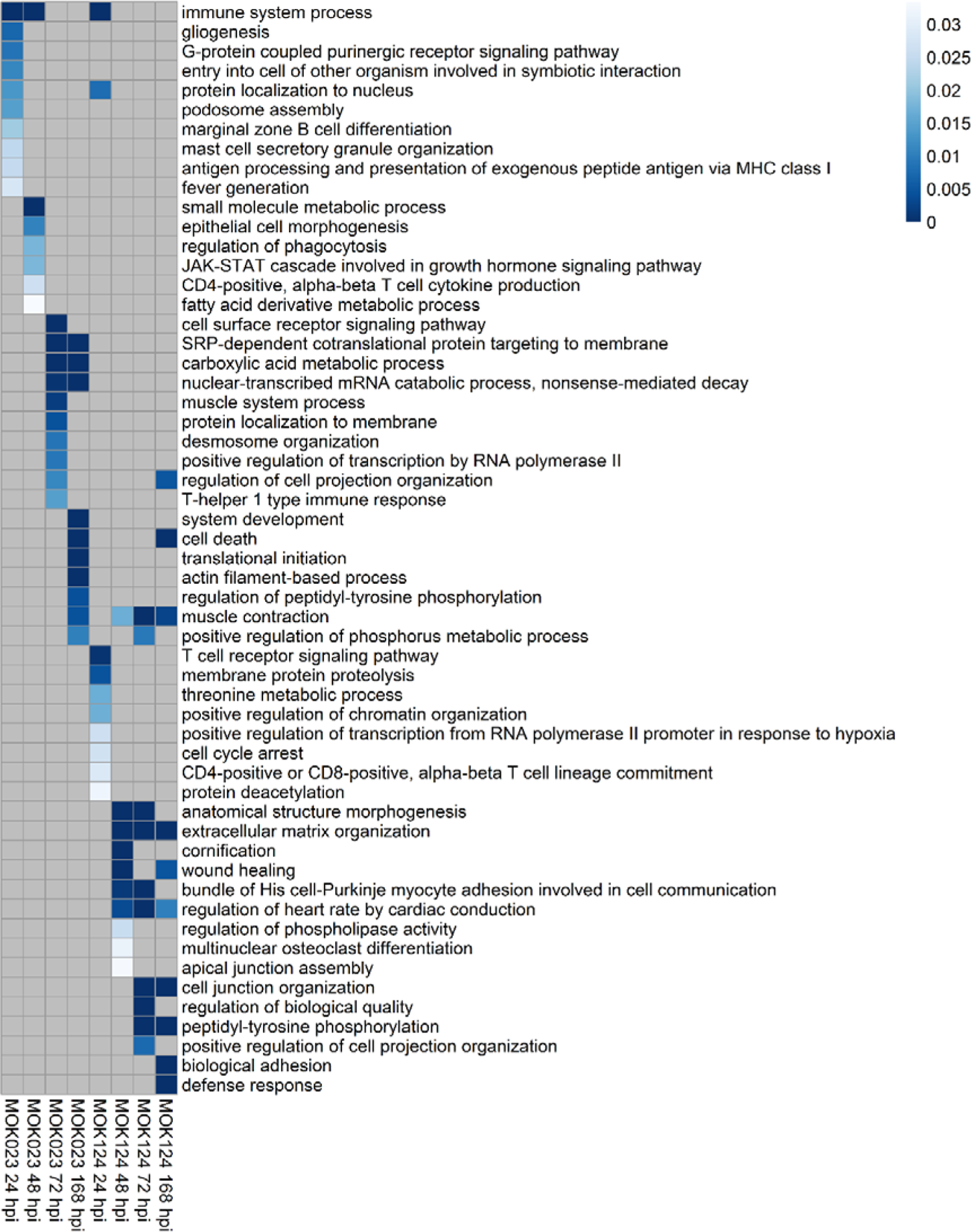
GO biological processes in response to infection with S. aureus MOK023 or MOK124 over time. Top 10 most significant (P < 0.05) and unique (filtered for shared genes) are shown. Colour scale represents P value, grey squares represent non-significance. hpi - hours post infection.

Immune system processes were overrepresented in the MOK023 group at 24 and 48 hpi, while they were only significant in MOK124 group at 24 hpi. While the GO terms in the MOK023 group varied across time points, a small number were consistently overrepresented in the MOK124 group between 48 and 168 hpi. These included extracellular matrix organisation, wound healing and cell junction organisation. Additionally, cell death was an enriched process in both groups at 168 hpi.

### Immune cell populations

A significantly higher proportion of neutrophils was observed in milk from cows in the MOK124 group than in the MOK023 group (30). In order to assess whether immune cell proportions in the milk somatic cell samples could be inferred using transcriptomic data, a cell population analysis based on gene expression of cell markers was performed using quanTIseq (43). The cell proportions at 0 hpi were similar between the two groups, with most cells classified as “other” likely representing mammary epithelial cells (Fig. 8A). Genes for epithelial cell markers such as cytokeratins 14, 18 and 19 (KRT14, KRT18, KRT19), E-cadherin (CDH1) and epithelial cell adhesion molecule (EPCAM) (44, 45) showed the highest expression at 0 hpi and decreased post-infection (Fig. 8B). Immune cell populations predominated at later time points and immune cell proportions differed between the two groups (Fig. 8A). At 24 hpi, neutrophils represented the largest proportion of milk somatic cells in both MOK023 and MOK124 infected animals; however, in the MOK124 group neutrophils represented a greater proportion of somatic cells than in the MOK023 group. A major difference in cell proportions was observed between 48 and 168 hpi. In the MOK023 group, the largest proportion of cells between 48 and 168 hpi were M1 macrophages, whereas in the MOK124 group neutrophils were the predominant cell type. A slightly larger proportion of M2 macrophages could also be observed at each time point post-infection in the MOK124 group when compared to the MOK023 group. M2 macrophages seemed to increase in proportion in both groups over time.

**Fig. 8.**
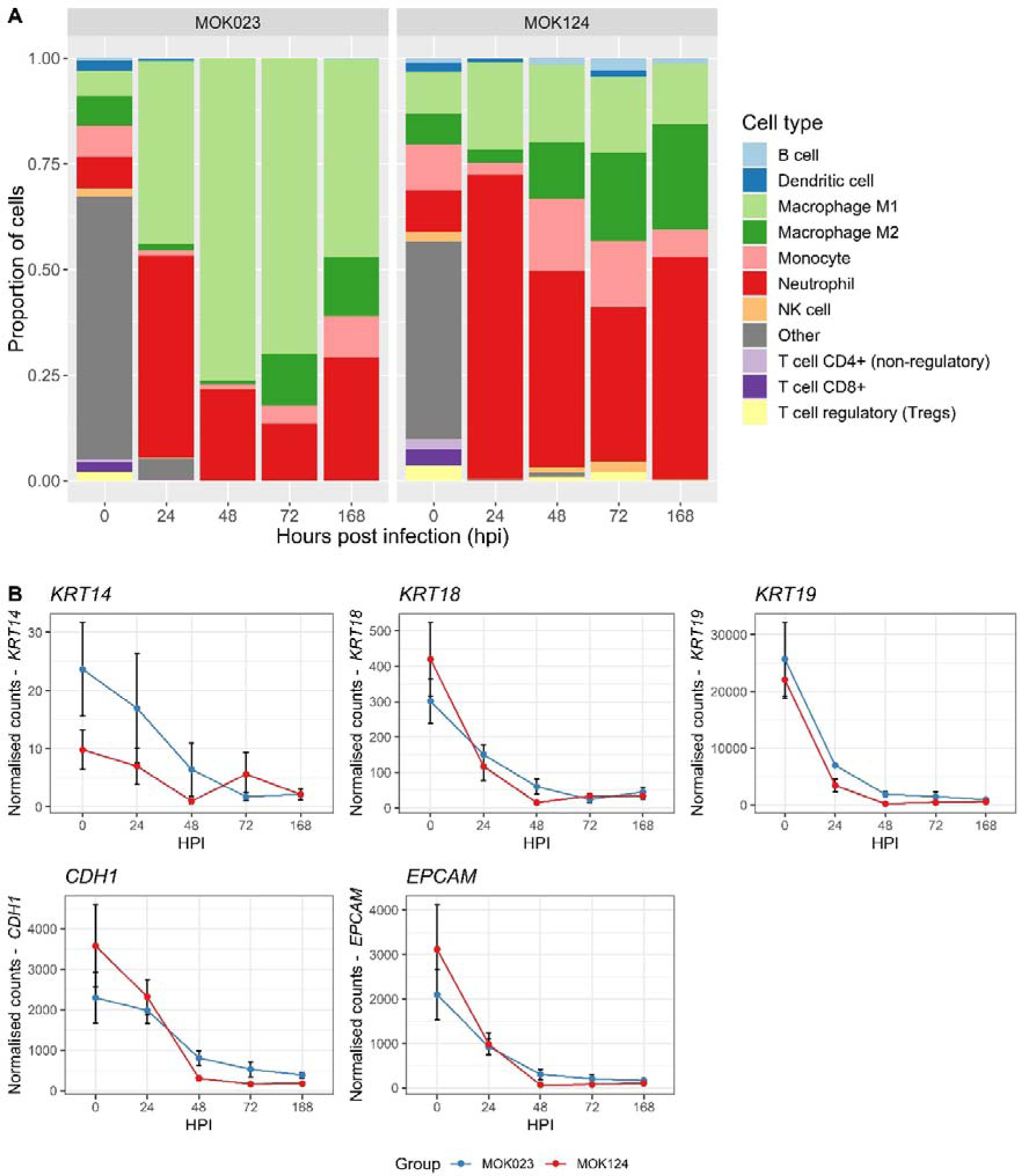
Cell composition analysis. a Changes in cellular composition in milk somatic cells over time illustrated by cell proportions. B Expression of epithelial cell marker genes in milk somatic cells over time in response to infection with MOK023 (blue) or MOK124 (red). KRT14-cytokeratin 14, KRT18 - cytokeratin 18, KRT19 - cytokeratin 19, CDH1 - E-cadherin, EPCAM - epithelial cell adhesion molecule, HPI - hours post infection.

### Milk secretion of chemokines and cytokines

Cytokine concentration was measured in milk at 0, 7, 24, 48, 72, 168, 336 and 504 hpi in order to reflect both the acute and chronic response to infection. There were no differences in the concentration of any of the examined cytokines or chemokines between the MOK023 and MOK124 groups prior to intramammary challenge or after 168 hpi. Protein concentrations increased post-infection in both groups with a peak expression at 168 hpi, except for TNFα and IL-8 which demonstrated a sharp increase at 24 hpi. A significant group x time interaction (P < 0.05) was observed for IL-1α, IL-1β, IL-1RA, and IL-6, which are typical pro-inflammatory cytokines, but also IL-2, IL-4, IL-10 and CCL3 (Fig. 9). In the case of all the above cytokines, MOK124 caused significantly higher protein secretion at 168 hpi, which is also a time point at which the difference in milk yield was low.

**Fig. 9.**
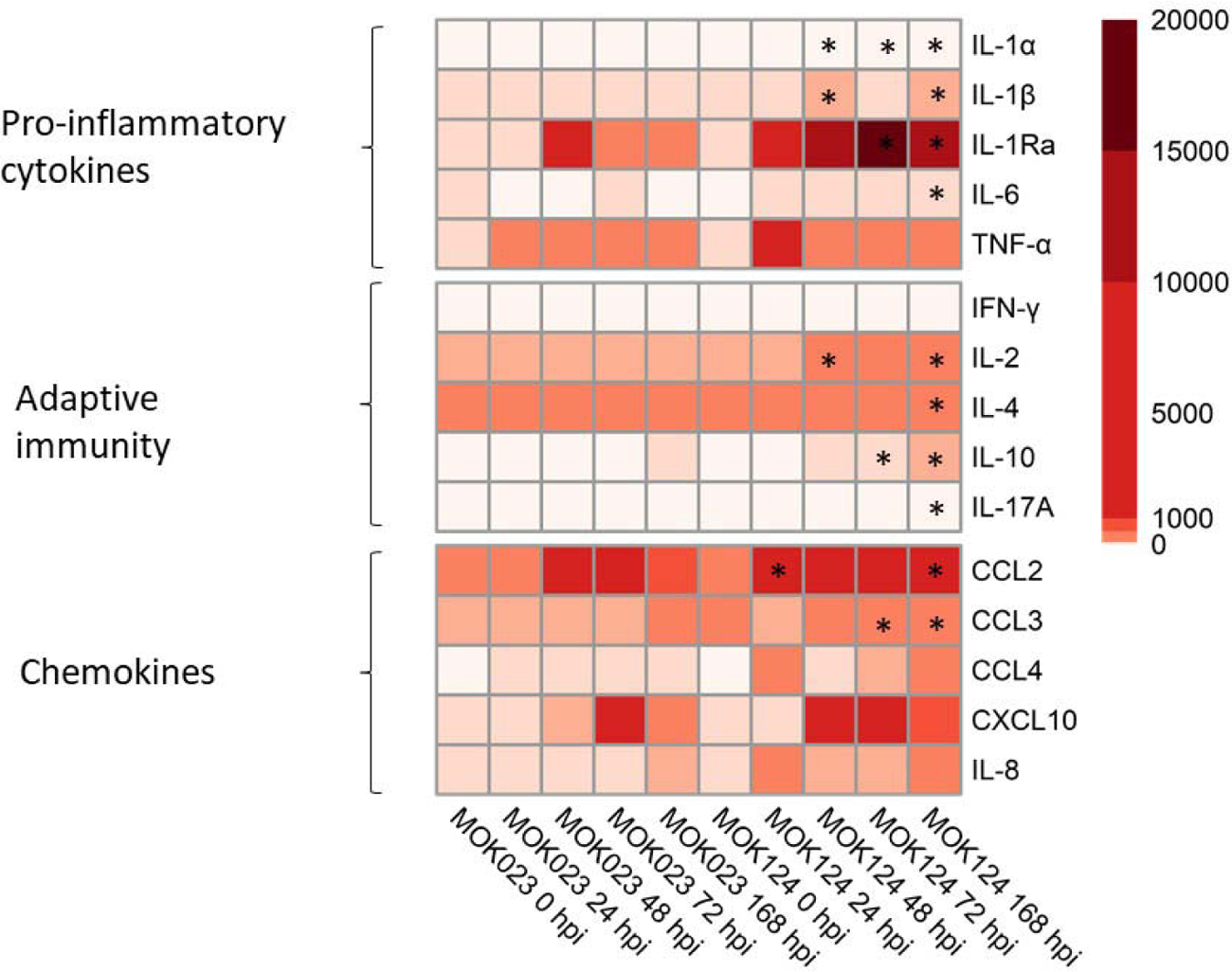
Concentration of selected immune markers in milk. The immune markers are divided into: pro-inflammatory cytokines: IL-1α, IL-1β, IL-1RA, IL-6, TNFa; cytokines involved in adaptive immunity: IFNγ, IL-2, IL-4, IL-10, IL-17A; and chemokines: CCL2, CCL3, CCL4, CXCL10, IL-8. Colour scale represents mean fluorescence intensity (MFI). Significant time points where cytokine concentration in MOK124 group was higher than in the MOK023 group are marked with *.

## Discussion

In this study, the transcriptional response of milk somatic cells to two genotypically distinct strains, representing two important globally distributed bovine-adapted lineages of S. aureus, was assessed. Differences in the response to the 2 strains were found in gene expression patterns and in overrepresented pathways. An initial immune response to both strains was evident but the transcriptional immune response was sustained in the MOK023 group between 24 and 72 hpi, while immune genes were only significantly upregulated in the MOK124 group at 24 hpi. This may indicate a difference in disease progression or a response to different virulence strategies employed by the two strains. The expression of genetic markers of various immune cells also differed in response to each strain, indicating differing immune cell proportions among somatic cells recruited in response to each strain, a result that was confirmed by a differential milk somatic cell count using microscopy (30). More neutrophils were recruited in response to MOK124 from 48 hpi and there were proportionally more M1 macrophages in the MOK023 group.

Exploration of normalised gene expression data yielded an interesting insight into the dynamics of the transcriptional response to the two infecting strains, with a strain-specific transcriptional response evident by 48 hpi. At this time point, gene expression in the MOK023 group was similar to that at 24 hpi while in the MOK124 group gene expression was more similar to that at 72 and 168 hpi. Examination of normalised counts of NFKB1 and other key genes suggested a difference in the pro-inflammatory response, with a more rapid and higher response to MOK124, which later subsided; in contrast, more sustained upregulation of immune gene expression was evident in response to MOK023. The sustained transcriptional upregulation of immune genes in the MOK023 group was supported by pathway analysis – the overrepresentation of both the NF-κB and TNF signalling pathways was sustained for longer after infection with MOK023. This difference in the transcriptional response may be related to a rapid switch from activation of the immune response to a disruption in cellular proliferation and the integrity of the epithelial cell layer in the MOK124 group. The 168 h time point could represent the end of the acute innate immune response in both groups, since immune pathways were not significantly enriched at this time point and few genes involved in the NF-kB and TNF signalling pathways were upregulated in either group.

The transcriptional response observed in this study supported several findings of the corresponding in vivo infection, such as the larger reduction in milk yield, higher inflammatory response at 24 hpi and higher SCC in the MOK124 group, as well as the increase in the proportion of neutrophils in milk observed in the MOK124 group. The downregulation of expression of milk production genes in the MOK124 group coincided with a significant reduction in milk yield, which was observed in response to MOK124 between 24 and 144 hpi when compared to MOK023 (30). This downregulation also indicates that infection with MOK124 caused a greater disruption to the epithelial layer of the mammary gland, possibly due to epithelial cell dysfunction and sloughing, when compared to MOK023 infection. Overall, the cellular composition analysis indicated that the strain-specific response was mediated by different cell populations. M1 macrophages were mainly involved in the response to MOK023, while the response to MOK124 led to a higher neutrophil recruitment. There was no significant difference between the MOK023 and MOK124 groups in the proportion of large mononuclear cells by microscopy; however, large mononuclear cells would have been comprised of M1 and M2 macrophages as well as monocytes, and shifts in those respective populations could be observed when looking at expression of cellular markers. The transcriptomic data therefore has the potential to provide more detail on the proportions of different cell populations in milk, especially in species where antibodies to cell markers are not available for the fine characterization of these cell subpopulations.

A larger proportion of M2 macrophages in somatic cells from the MOK124 group may indicate a trend towards tissue repair. M2 macrophages synthesise polyamines and proline which stimulate cell growth, collagen formation and tissue repair (46). M2 macrophages are also capable of producing ECM components (47) and may have contributed to the ECM related pathways which were overrepresented in response to MOK124. Therefore, it is possible that MOK124 creates high epithelial dysfunction, which exposes the stroma to bacteria and bacterial products and induces a strong inflammation that the organism attempts to repair. This leads to a higher recruitment of neutrophils and more plasma leakage due to failure of the epithelium to function as a barrier.

Pathway analysis revealed that MOK124 elicited a strain-specific response indicative of tissue injury, as illustrated by the downregulation of Hippo signalling, ECM receptor interaction and tight junction pathways between 48 and 168 hpi. Moreover, MOK124-specific GO terms identified at those time points, such as wound healing and ECM organisation, corresponded to the KEGG pathways identified. These pathways are well studied in humans. The key genes in the Hippo signalling pathway are YAP1 and WWTR which encode transcription factors YAP and TAZ. A successful activation of this pathway depends on translocation of the YAP/TAZ proteins to the nucleus, followed by binding of the transcription factors to a TEAD protein. The YAP/TAZ-TEAD complex causes activation of transcriptional responses which result in cell proliferation by controlling the expression of genes directly involved in the cell cycle and by upregulation of anti-apoptotic genes (40). The genes involved in the Hippo signalling pathway were downregulated, which could indicate a decrease in cell proliferation. However, protein interactions and localisation in the cell would need to be studied in order to confirm this hypothesis.

The ECM receptor interaction pathway is involved in maintenance of cell shape and the interaction of epithelial cells with the basement membrane. Within the ECM receptor interaction pathway, several integrin genes were upregulated in response to MOK124 at 48 hpi (ITGA5, ITGA9, ITGB7, ITGA2B). The integrin proteins mediate cell to matrix and cell to cell adhesion (48). ITGA5 encodes the light and heavy chains that comprise the α5 subunit. This subunit associates with the β1 subunit to form a fibronectin and fibrinogen receptor. Both fibronectin and fibrinogen can mediate adhesion of S. aureus to host cells (49), and increase of fibronectin levels in whey of Holstein-Friesian cows is associated with mastitis progression (50). The integrin α9 subunit encoded by the ITGA9 gene, when bound to the β1 chain, forms an integrin that is a receptor for vascular cell adhesion molecule 1 (VCAM1), cytotactin and osteopontin, all of which are involved in immune response. ITGB7 encodes integrin subunit β7 which forms dimers with an α4 chain or an αE chain and plays a role in leukocyte adhesion (48). These two upregulated integrin genes could indicate leukocyte adhesion and migration was taking place in MOK124 infected animals at 48 hpi. Integrins can also act with cytoskeleton proteins from the tight junction pathway and clustering of integrins increases cellular tension, which can lead to the activation of the Hippo signalling pathway (48).

Tight junctions form barriers between cells to the flux of ions and molecules. Pore and leak pathways can be differentiated within the action of the tight junction, and while claudins are involved in the pore pathway, zonula occludens 1 (ZO-1) and occludins are involved in the leak pathway (51). In the MOK124 group, claudin (CCLN), occludin (OCLN), Marvel (MARVELD3) and immunoglobulin superfamily member (IGSF5) genes were downregulated as part of the tight junction pathway, which suggests a decrease in cell to cell interaction and cell detachment. TNF-α has been shown to induce barrier loss in epithelial cells during Crohn’s disease in vitro and in mice. The loss of integrity of intestinal epithelium in those studies was mediated by TNF-induced occludin internalisation (51). Therefore, upregulation of the TNF pathway in the MOK124 group at 24 hpi could have contributed to loss of cell integrity at later time points. On the other hand, dysregulation of cytoskeleton genes in the tight junction pathway could be indicative of S. aureus internalisation into host cells. Modification of the actin cytoskeleton in cells infected by S. aureus has previously been described (28) and was attributed to host cell internalisation. A CC151 strain, RF122, has been hypothesised to be adapted to an intracellular niche (19). In vitro, strain MOK023 was a significantly higher internaliser in bMEC than strain MOK124 (34). The cytoskeleton reorganisation in response to MOK124 is therefore more likely to be related to cell damage, loss of cellular structure and an attempt to repair the tissue rather than due to host cell internalisation. The presence of clinical signs in MOK124 challenged animals (30) also confirms that cell damage likely took place and a healing response would be necessary to rebuild the damaged tissue. The ability of MOK124 to cause tissue damage may be associated with the toxin encoding genes present in this lineage. MOK124 encodes enterotoxin genes seg, sei, selm, seln, seln, selo, egc, selu and ORFCM14 as well as the leukocidins lukF-PV and lukM, none of which are present in MOK023 (17). Staphylococcal enterotoxin genes seg, sei, selm and sei as well as agrII (which is the agr type of MOK124) have recently been found to be associated with a higher likelihood of causing clinical mastitis in a collection of European S. aureus strains isolated from natural infections (18). While LukM’ was not found in milk of MOK124 infected cows in this study, the other toxins encoded by this strain could play a role in the virulence of this strain. Other CC151 strains have been found to produce low amounts of LukM’ in previous studies (52), therefore the mechanism of neutrophil killing by CC151 strains remains to be determined. Since immune genes tended to be upregulated for longer in response to MOK023 than in response to MOK124, the greater influx of neutrophils in response to MOK124 is unlikely entirely due to immune signalling by milk somatic cells. Instead, a larger influx of neutrophils could be related to cellular and tissue damage.

Although MOK023 caused immune gene induction both in bMEC in vitro and in milk somatic cells in vivo (30, 34), immune signalling from bMEC was followed by a temporary recruitment of neutrophils at 24 hpi, and M1 macrophages became the predominant cell type from 48 hpi. Since macrophages are normally present in a healthy mammary gland, while neutrophils tend to be recruited to a site of intramammary infection (53), the difference in cellular composition in milk post-infection in response to the two strains may indicate that MOK023 subverted the immune response which failed to recruit neutrophils to a significant extent. This is also supported by the fact that even though immune gene expression was sustained in response to MOK023, bacterial load increased in the MOK023 infected animals, to reach its maximum at 168 hpi. Lack of inflammation and relatively low neutrophil numbers in response to MOK023 could be due to host cell internalisation. It was demonstrated that MOK023 can internalise into bMEC in vitro (34). It has also been reported that human-associated S. aureus can internalise and survive within human (54) and bovine macrophages (55). S. aureus SH1000 (CC8) internalised within murine or human macrophages can exhibit a persister phenotype and evade antimicrobial treatment (56), and strain ATCC 29213 (CC5; (57)) was found to promote its survival in bovine macrophages by blocking autophagic flux (55). Internalisation of MOK023 into MEC and/or macrophages could aid in the evasion of immune response. Interestingly, among significantly DE genes, LAMP1 was found to be overexpressed solely in response to MOK023 (Supplementary File 2). This gene has previously been associated with S. aureus survival inside a mature phagosome in macrophage cell lines (58, 59). Upregulation of the tuberculosis pathway in the MOK023 group further supports the hypothesis that MOK023 may internalise into host cell macrophages and be immune evasive. The tuberculosis pathway represents a process where tubercle bacilli enter macrophages in the lung, and following host cell internalisation they interfere with phagosomal maturation, antigen presentation, apoptosis and host bactericidal responses to establish persistent or latent infection (60). Persistence of infection of MOK023 was illustrated by a consistent bacterial load in infected animals over 30 days; however, more research is needed to determine the precise mechanisms by which this strain persists within the host. Notably, a recent study associated high internalisation into bMEC, biofilm formation and low cytotoxicity with high within-herd prevalence of a S. aureus CC9 strain (61).

The transcriptomic response to S. aureus mastitis in dairy cows has been studied previously with samples taken at single time points: 3 hpi (28, 32), 24 hpi (33) and 48 hpi (62). Udder biopsies were used in those studies to analyse gene expression in response to a S. aureus IMI, and single S. aureus strains were used. This study characterises later events occurring in IMI, and represents a response of different cell populations, as it would be expected that an udder biopsy would be mostly comprised of bMEC. Transcriptomic analysis of S. aureus mastitis in naturally infected dairy cows has also been performed on post-mortem mammary tissue and peripheral blood leukocytes (63, 64), with the extent of time post-infection being unknown, and the infecting strains not identified. This is the first transcriptomics study of S. aureus mediated IMI where the response to two different strains of S. aureus are compared, with full strain genotype information available. Genotyping of bacterial strains used, use of multiple strains of S. aureus as well as examination of the cellular populations of studied samples should be considered in future studies. All of the above can ensure better understanding of strain-specific responses and an improved ability to compare different studies of IMI.

The concentration of selected cytokines and chemokines in milk throughout the course of infection confirmed and extended the finding that MOK124 induced higher IL-1β and IL-8 concentrations in milk. A discrepancy was found between the gene expression results and the cytokine profiles since cytokine and chemokine protein levels remained elevated in response to MOK124 at later time points, while most of the cytokine genes were not DE post 24 hours in this group. While correlation between transcriptomic and proteomic data is often poor (65–67) due to protein turnover, RNA degradation or post-translational modifications, there could be other reasons for the discrepancy in this study. Firstly, the protein levels were measured in milk, while the transcript levels were measured in milk somatic cells. The mammary epithelial cells lining the bovine mammary gland could be contributing to cytokine secretion into milk, and would not be captured by the RNA sequencing. Therefore, mammary epithelial cells and other cells of the epithelial lining such as resident macrophages and lymphocytes could be major contributors to cytokine and chemokine levels in the milk. Secondly, the number and the type of milk cells likely influenced absolute protein concentration with higher neutrophil counts in MOK124 infected cows. Finally, a systemic response could have contributed to increased milk cytokine levels as exudation of plasma was likely in the MOK124 group with higher milk Ig concentrations, but not evident in the MOK023 group (30). The milk cytokine data highlights the need to implement proteomic studies alongside transcriptomic studies to better understand the molecular mechanisms of host immune response. A study where each milk somatic cell type was assessed separately during an in vivo trial, along with udder biopsies or post-mortem examinations at one pre-defined time point, would be beneficial to further understanding of bovine immune response to S. aureus. Furthermore, the pattern of cytokine expression measured in vitro on MEC culture infected with either strain (34) does not correlate with the secretion profile detected in milk upon mammary gland infection of dairy cows, showing that bacteria-MEC interactions do not summarize the complex mechanisms occurring during mastitis and may not be predictive of the pathogenicity of S. aureus strains.

## Conclusions

This study confirms the ability of bovine S. aureus strains to cause a strain-specific immune response during IMI, with each strain resulting in a unique signature of infection in terms of the number and type of immune cells recruited, the biological function of the recruited cells and the resulting cytokine and chemokine concentration in milk. MOK124 caused more inflammation and udder damage, as illustrated by the disruption of tight junction and Hippo signalling pathways between 48 and 168 hpi, coupled with clinical signs observed in the infected animals. On the other hand, MOK023 caused less inflammation and may be able to limit neutrophil recruitment and hence persist in the mammary gland, as suggested by the sustained immune gene expression but low increase in SCC, sustained bacterial load and mild clinical signs in MOK023 infected cows. Since differences in early transcriptional response between the two strains were observed, followed by either a severe or a mild, persistent infection, it is possible that the magnitude of this early response may be crucial in determination of the course of staphylococcal IMI. The response to each strain also involved recruitment of specific immune cells, with M1 macrophages primarily recruited in response to MOK023, while neutrophils were the main immune cell type recruited in response to MOK124. This indicates that different cell types as well as the transcriptional activity of each cell type may influence gene expression differences between groups when analysing mixed cell population tissues such as blood or milk. Further studies are necessary to further understand the molecular mechanisms utilised by these strains to cause inflammation or evade the immune response.

## Methods

### Study design

The aim of this study was to elucidate the molecular mechanisms of the host immune response in vivo by utilising a transcriptomic approach. Fourteen first lactation Holstein-Friesian cows were purchased from two commercial farms which were leptospirosis, salmonellosis and Johne’s disease free. Animals were selected based on composite milk SCC < 100,000 cells/ml. Milk bacteriology was performed prior to challenge to confirm lack of contaminating organisms or mastitis pathogens. Cows were infected with approximately 500 cfu of S. aureus strain MOK023 (ST3170, CC97) or MOK124 (ST151, CC151) in 400 µl of PBS as described previously (30). Animals in which a S. aureus infection was not established (two cows challenged with MOK023 and one cow challenged with MOK124) were excluded from the study. Therefore, the MOK023 infected group included 5 animals, while the MOK124 infected group included 6 animals. Due to severe clinical mastitis developed by 2 cows in the MOK124 group, these animals were treated with antibiotic and milk samples for transcriptomic analysis were not taken after antibiotic treatment: cow 1776 was removed from the trial at 31 hpi, while cow 1920 was removed at 120 hpi. A further sample from the MOK124 group at 48 hpi was removed from the analysis for quality control purposes as a sample from an uninfected quarter was mistakenly sequenced instead of the sample from the infected quarter. The number of animals in each experimental group at each time point is presented in Table 2.

**Table 2.** The numbers of animals (N) included in the study at each time point.

| Hours post-infection (hpi) | N, MOK023 group | N, MOK124 group |
| --- | --- | --- |
| 0 | 5 | 6 |
| 24 | 5 | 6 |
| 48 | 5 | 4 |
| 72 | 5 | 5 |
| 168 | 5 | 4 |

### RNA extraction from milk somatic cells

Milk from the infected quarter was collected into a sterile universal tube (Sarstedt, Nümbrecht, Germany). In order to obtain a sufficient quantity of RNA, 100 ml of milk was used for somatic cell isolation. Milk was centrifuged at 800 x g, at 15 °C for 10 min. The fat layer and supernatant were removed and tubes were placed upside down on tissue paper to drain for 10 min. Cell pellets were subsequently resuspended in 900 μl of Qiazol (Qiagen, Manchester, UK) and vortexed for 20 s to lyse the cells. Lysed samples were stored in Qiazol at −20 °C until RNA extraction. Samples were rapidly thawed at 37 °C and maintained at room temperature (RT) for the duration of the extraction. RNA was extracted using an RNeasy Plus Universal kit (Qiagen, Manchester, UK) as per the manufacturer’s instructions. RNA was eluted in 30 μl of RNase-free water (Sigma Aldrich Ireland Ltd., Wicklow, Ireland).

RNA quality was assessed using 2100 Bioanalyser RNA Nano chips (Agilent Technologies Ireland Limited, Cork, Ireland) as well as with a NanoDrop™ 1000 spectrophotometer (Thermo Fisher Scientific, Waltham, MA, USA). RNA quantity was determined with a Qubit RNA broad range assay (Invitrogen: Bio-Sciences, Dun Laoghaire, Ireland). Sample RIN values ranged between 5.9 and 9.5 with a mean RIN value of 7.82 (Supplementary Table S1), and concentrations ranged from 4.1 – 930 ng/µl (mean concentration of 154 ng/μl).

### Library preparation and sequencing

RNA was sent on dry ice to Macrogen (South Korea) for library preparation and sequencing. RNA quality was confirmed on arrival using a TapeStation HSRNA Screen Tape and analysed on a 2200 TapeStation (Agilent Technologies Korea Ltd, Seoul, Korea). Library preparation was performed with 100 ng of RNA using a TruSeq Stranded mRNA LT Sample Prep Kit (Illumina, San Diego, USA) according to the manufacturer’s instructions (Part #15031047 Rev. E). Library size was verified with a 2100 Bioanalyzer using a DNA 1000 chip (Agilent, Korea) and library quantity was determined with qPCR according to the Illumina qPCR Quantification Protocol Guide. The mean library fragment size was 348 bp while the mean library concentration was 92.63 nM. Libraries were diluted and pooled with 300 pM of each library and sequenced (100 bp, paired-end) on a NovaSeq (Illumina, USA). Samples were divided across two sequencing runs.

### Quality control, assembly and DE gene analysis

All the bioinformatics pipeline bash, Perl and R scripts used for computational analyses were deposited in a GitHub repository at https://github.com/DagmaraNiedziela/RNAseq_Saureus_cattle_infection. RStudio v1.1.463 and R v3.5.2 were used for the analyses. Sequence quality was assessed using FastQC v11.5 (68). Trimming was performed using fastp v12.1 (69) with default settings as well as enabling base correction for paired end data and enabling overrepresented sequence analysis; on average 0.5 % of the reads were trimmed per sample. Trimmed fastq files were re-checked with FastQC to confirm sequence read quality. Trimmed sequences were mapped to the bovine genome (UMD3.1) using the STAR aligner v2.5.2 (70), with gene counts generated simultaneously with the STAR software using the UMD3.1 v93 of the B. taurus reference genome and annotation. Genes with zero read counts across all samples as well as non-protein coding genes were removed in a filtering step (71). DeSeq2 v1.18.1 (72) was then used to visualise gene expression data and perform differentially expressed (DE) gene analysis. Data was normalised with a vst function and visualised with a principal component analysis (PCA) plot of the samples based on the 500 genes with the highest inter-sample variation. DE gene lists were generated using a negative binomial generalized linear model. Wald tests were performed to obtain DE genes separately for each group at each time point using time 0 as a reference, with P values adjusted for multiple comparisons using a Benjamini and Hochberg (B-H) method. DE genes with an adjusted P value < 0.05 and a log_2_ fold change threshold of 1 were used for further DE gene data exploration and pathway analysis.

### Pathway and gene correlation analysis

Significant (P adj < 0.05) and highly overexpressed (|log_2_ fold change|> 1) DE genes were converted to their 1 to 1 human orthologs using BioMart, ordered by adjusted P value and analysed for enrichment of Gene Ontology (GO) terms and Kyoto Encyclopedia of Genes and Genomes (KEGG) pathways using gProfiler R package v0.6.6 (73). When a group of DE genes matched to multiple pathways, the pathway that was represented by the highest number of DE genes was selected in a filtering step.

### Cell composition analysis

Transcripts per million (TPMs) were generated from raw gene counts. Bovine Gene IDs converted to 1 to 1 human orthologs were used for the analysis. To estimate the proportion of the different immune cell populations in each somatic cell sample, a quanTIseq algorithm which is based on expression of cell specific surface marker genes was used (43).

### Quantification of a panel of cytokines and chemokines in milk

Milk was defatted by centrifugation at 800 x g, 15 °C for 10 min. The top fat layer was discarded and the skim milk supernatant was stored in 1 ml aliquots at −20 °C until use. Prior to protein quantification milk was diluted 1:2 in assay buffer (Merck-Millipore, Burlington, USA). Concentrations for 15 cytokines (IL-1α, IL-1β, IL-1RA, IL-2, IL-4, IL-6, IL-17a, IFNγ, CCL-2, CCL-3, CCL-4, CXCL8, CXCL10, and TNFα) were determined using a custom bovine cytokines MilliPlex xMAP assay (Merck Millipore, USA). Data were recorded on a MAGPIX flow cytometer using Xponent software (Luminex, Austin, USA).

### LukM ELISA

ELISA to detect the LukM subunit of the LukMF’ Staphylococcal toxin in milk samples was performed as described previously (37). Briefly, anti-LukM polyclonal bovine IgG (32 µg/ml) was used to coat the wells of an ELISA plate at 4 °C overnight. The plate was emptied and washed twice with 300 µl of PBS-T (DBPS: Gibco, UK; 0.05% Tween-20: Sigma-Aldrich, St. Louis, MO), blocked with 200 µl of 4 % skimmed milk powder (Marvel, DSDelta Ltd) in PBS-T for 30 min with shaking and washed once with 300 µl PBS-T. A 100 µl aliquot of each sample (skimmed milk heated to 95 °C for 10 minutes to inactivate any existing antibodies against LukM) or standard (recombinant LukM; kindly provided by Utrecht University) was added to the wells and incubated for 60 min at room temperature (RT) with shaking, followed by 2 washes with 300 µl PBS-T. Mouse anti-LukM monoclonal antibody (KoMa43, Podiceps, The Netherlands) was diluted to 3 µg/ml in 1 % skimmed milk-PBS-T and 50 µl was added to each well. The plate was incubated for 60 min at RT, with shaking, and washed 3 times. Polyclonal HRP conjugated goat anti-mouse IgG (BioLegend, London, UK) was diluted 1 in 1000 and 100 µl was added to each well. The plate was incubated for 60 min at RT, with shaking, and washed 3 times. TMB (100 µl; Thermo Fisher Scientific, Waltham, USA) was subsequently added and incubated in the dark at RT for 15 min, followed by addition of 100 µl of the stop solution (0.05 M H_2_SO_4;_ Thermo Fisher Scientific, Waltham, USA). Absorbance was read at 450 nm using a microplate reader (xMark, Bio-Rad, Watford, UK).

## Statistical analysis

Data generated from cytokine and LukM’ protein quantification was analysed with a repeated measures ANOVA using a MIXED® procedure in SAS (SAS Version 9.4, SAS Institute, Cary NC, USA) to compare strain-dependent differences. Strain, time and the strain x time interaction were included in the model, with time as a repeated measure. Time 0 measurements for each cytokine were included as a covariate, and all variable interactions were explored. Log_10_ transformations were used for all cytokine variables to satisfy the distributional requirements of ANOVA. Correlations among repeated measurements across time between strains were modelled using the compound symmetry covariance structure for each parameter analysed. Tukey’s post-hoc multiple comparison test was applied as appropriate.

## Declarations

### Ethics approval

All procedures involving animals were performed in accordance with the relevant guidelines and regulations of the European Directive 2010/63/EU and received approval from the University College Dublin Animal Research Ethics Committee (protocol number Leonard AREC 16 44), and a project authorisation from the Health Product Regulatory Authority (HPRA) (project authorisation number AE18982/P108). The study was carried out in compliance with the ARRIVE guidelines and has been described in detail previously (Niedziela DA, et al. 2020. J Dairy Sci. 103(9):8453-66).

## Consent for publication

Not applicable.

## Availability of data and materials

Raw fastq files used during the current study are available in the European Nucleotide Archive (ENA) repository with a project number PRJEB43443. Code used to analyse the data is available at: https://github.com/DagmaraNiedziela/RNAseq_Saureus_cattle_infection Any other data from the current study are available from the corresponding author on reasonable request.

## Author contributions

DAN optimised and conducted the laboratory-based work, performed bioinformatics analysis of the RNAseq data and drafted the manuscript. OMK conceived the initial idea of the study and participated in its design and coordination. FCL conceived the initial idea of the study and participated in its design and coordination. PC provided mentoring and supervision during the analysis of RNAseq data and performed the cell composition analysis. GF conducted/supervised protein laboratory-based work and statistical analysis of protein data. All authors helped to draft the manuscript and read and approved the final manuscript.

## Acknowledgements

We would like to acknowledge the farm staff in UCD Lyons Research Farm, particularly Eddie Jordan, Michael Clarke and Joseph Callanan for invaluable assistance with sample collection for this study. Thank you to Dr Matthew McCabe (Teagasc) and Dr. Elaine Kenny (ELDA Biotech) for advice on RNA extraction. The LukM’ ELISA was performed by Margaret Murray in Teagasc Grange. Antibodies and protocol for this ELISA were generously provided by Manouk Vrieling, Kok van Kessel and Lindert Benedictus (Utrecht University).

## Abbreviations

bMEC: bovine mammary epithelial cells

CC: clonal complex

DE: differentially expressed

ECM: extracellular matrix

hpi: hours post-infection

GO: Gene Ontology

IMI: intramammary infection

KEGG: Kyoto Encyclopedia of Genes and Genomes

PCA: principal component analysis

RT: room temperature

ST: sequence type. Gene symbol abbreviations – Supplementary Table S2.

